# MASH Native: A Unified Solution for Native Top-Down Proteomics Data Processing

**DOI:** 10.1101/2023.01.02.522513

**Authors:** Eli J. Larson, Melissa R. Pergande, Michelle E. Moss, Kalina J. Rossler, R. Kent Wenger, Boris Krichel, Harini Josyer, Jake A. Melby, David S. Roberts, Kyndalanne Pike, Zhuoxin Shi, Hsin-Ju Chan, Bridget Knight, Holden T. Rogers, Kyle A. Brown, Irene M. Ong, Kyowon Jeong, Michael Marty, Sean J. McIlwain, Ying Ge

## Abstract

Native top-down proteomics (nTDP) integrates native mass spectrometry (nMS) with top-down proteomics (TDP) to provide comprehensive analysis of protein complexes together with proteoform identification and characterization. Despite significant advances in nMS and TDP software developments, a unified and user-friendly software package for analysis of nTDP data remains lacking. Herein, we have developed MASH Native to provide a unified solution for nTDP to process complex datasets with database searching capabilities in a user-friendly interface. MASH Native supports various data formats and incorporates multiple options for deconvolution, database searching, and spectral summing to provide a one-stop shop for characterizing both native protein complexes and proteoforms. The MASH Native app, video tutorials, written tutorials and additional documentation are freely available for download at https://labs.wisc.edu/gelab/MASH_Explorer/MASHNativeSoftware.php. All data files shown in user tutorials are included with the MASH Native software in the download .zip file.

## Introduction

Native mass spectrometry (nMS) analyzes intact proteins and protein complexes under non-denaturing conditions to preserve their tertiary structure and non-covalent interactions in the gas phase, which has emerged as a powerful structural biology tool to define protein structure-function relationships (Loo, 1997; Sharon and Robinson, 2007; Leney and Heck, 2017; Keener *et al*., 2021; Karch *et al*., 2022). Native top-down proteomics (nTDP) integrates nMS with top-down proteomics (TDP) (Catherman *et al*., 2014; Toby *et al*., 2016; Chen *et al*., 2018; Melby *et al*., 2021), which enables structural characterization of protein complexes together with proteoform sequencing to locate non-covalent ligand binding sites, posttranslational modifications (PTMs), and mutations (Li *et al*., 2018; Zhou *et al*., 2020; Karch *et al*., 2022; Jooß *et al*., 2022). nTDP first measures intact proteins and protein complexes under non-denaturing conditions (MS1) then directly fragments proteins and protein complexes in the gas phase (MS2) to obtain primary sequence information from a single dissociation event (Li *et al*., 2018). Alternatively, nTDP may be implemented in the “complex-down” mode using two separate dissociation events: 1) dissociation of intact protein complexes (MS1) into protein subunits (MS2’) by low-energy collision-induced dissociation (CID) or surface induced dissociation (SID), and 2) fragmentation of subunits (MS3) by tandem mass spectrometry techniques such as high-energy CID, electron capture dissociation (ECD), electron transfer dissociation (ETD) or ultraviolet photodissociation (UVPD) to provide primary sequence coverage and localize modifications (Skinner *et al*., 2018b; Stiving *et al*., 2019; Jooß *et al*., 2022).

Currently one of the major challenges in nTDP is the analysis of complex nTDP datasets which include both isotopically resolved and unresolved MS1 and MS2’ spectra as well as the complicated MS2 and MS3 data, and difficulties in database searching. Although multiple software packages have been developed for nMS of known proteins and complexes (Marty *et al*., 2015; Cleary *et al*., 2016, 2018; Reid *et al*., 2018), the lack of any MS2/MS3 fragmentation assignment and database searching prevent the identification of unknown proteins. Meanwhile, significant efforts have been allocated towards the development of software packages for denatured TDP with capability in analyzing complicated MS2/MS3 datasets with database search algorithms to identify unknown proteins (Sun *et al*., 2016; Kou *et al*., 2016; Fellers *et al*., 2015; Cai *et al*., 2016; Wu *et al*., 2020), but these denatured TDP software packages lack the capability to analyze the unresolved MS1/MS2’ that are characteristic of nMS data. Hence, there is a critical need for a universal software package to address this major challenge in nTDP that can process MS1, MS2, MS2’ and MS3 datasets with database search capabilities.

Herein, we introduce MASH Native (https://labs.wisc.edu/gelab/MASH_Explorer/MASHNativeSoftware.php), a unified solution for

nTDP which can process unresolved MS1 and MS2’ data together with isotopically resolved MS1, MS2, and MS3 deconvolution and database searching (**Figure 1**). MASH Native supports various nTDP applications in both targeted mode to characterize known proteins and discovery mode to identify unknown native proteins. It supports various MS file types with different vendor formats and integrates multiple deconvolution/search algorithms into one package. We detail the functions and features of MASH Native and provide examples of processing nTDP data to showcase its capabilities as a “one-stop shop” for nTDP.

**Figure 1.**
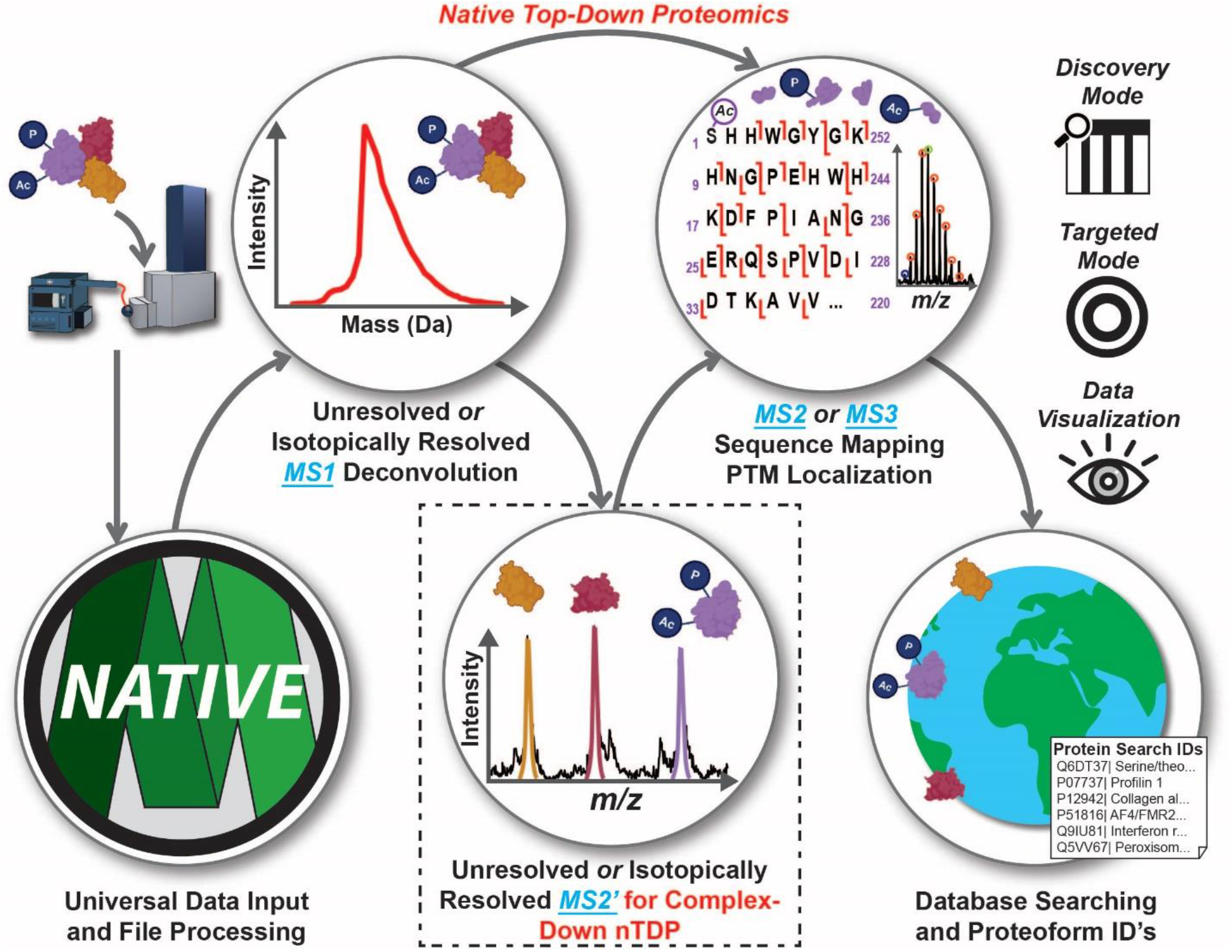
MASH Native provides a universal and comprehensive data processing software for a variety of nTDP analyses. MASH Native is capable of deconvoluting unresolved protein/protein complex (MS1) and released protein subunits (MS2’) spectra, deconvoluting isotopically resolved MS1, MS2’, MS2, and MS3 spectra, and performing database searches to identify unknown proteins. MASH Native can process nTDP data in both Discovery Mode and Targeted Mode approaches. It supports various MS file types and integrates multiple deconvolution/search algorithms into one package. MASH Native is a user-friendly software package capable of providing a “one-stop shop” for nTDP data processing.

## Results

The MASH Native user interface is a multithreaded windows desktop application written under a .NET framework environment in Visual Studio using the C# programming language (Wu *et al*., 2020). MASH Native provides universal MS file support through ProteoWizard’s file conversion engine, MSConvert (Chambers *et al*., 2012), and directly imports both vendor-specific MS file types (Thermo *.RAW, Bruker *.d/*.baf/*.ascii) and general file formats (*.mgf, *.mzML, *.mzXML).

MASH Native software can deconvolute both isotopically resolved and unresolved data at the MS1, MS2, and MS3 level and enables database searching of nTDP results (**Figure 1**). It can process nTDP, nMS, and complex-down proteomics data using multiple deconvolution and database search algorithms with flexible data output options (**Figure S1**). It also maintains the functions and capabilities previously developed for denaturing TDP so users can process both nTDP and TDP in the same software. To address challenges with low signal-to-noise (S/N) ratios of intact and fragment mass spectra, MASH Native includes a variety of spectral summing algorithms that may be applied prior to data processing workflows (**Figure S2 and S3**). To deconvolute unresolved MS1 spectra, MASH Native includes UniDec (Marty *et al*., 2015), a powerful deconvolution algorithm, to characterize both unresolved and isotopically resolved nMS data (**Figure S4**). Isotopically resolved spectral deconvolution can also be performed in MASH Native (**Figure S5**), including TopFD (Kou *et al*., 2016), MsDeconv (Liu *et al*., 2010), eTHRASH (Horn *et al*., 2000), and pParseTD (Yuan *et al*., 2012). Users may also import previously deconvoluted results from external deconvolution algorithms, such as FLASHDeconv (Jeong *et al*., 2020), ProMEX (Park *et al*., 2017) or Maximum Entropy (Ferrige *et al*., 1991). Deconvolution results can be combined into a single output table, allowing users view MS1, MS2, and MS3 results simultaneously and combine multiple deconvolution types to improve protein sequence coverage (Mcilwain *et al*., 2020). Results of deconvolution may be searched against a user-selected *.FASTA file or user-defined protein sequence with TopPIC (Kou *et al*., 2016), MS-Align+ (Liu *et al*., 2012), or pTop (Sun *et al*., 2016) to identify proteoforms in a complex mixture. Search results generated through MASH Native or from additional search tools such as MSPathFinderT (Park *et al*., 2017), may then be imported in MASH Native to view identifications, generate fragment ion maps, view fragment ions, and validate for all identified proteins and proteoforms.

To facilitate high-throughput data analysis, user-defined MASH Native processing workflows can be designed, saved, and queued to allow batch processing of data files using two different approaches: Discovery and Targeted Mode. Discovery Mode facilitates identification of unknown proteins though database searching, a critical processing feature absent from current nMS or native top-down software tools. This mode combines MS1 processing with isotopically resolved MS2 or MS3 deconvolution and database searching in a single workflow for nTDP datasets (**Figure S6**). To demonstrate MASH Native Discovery Mode for data processing, we accessed and reanalyzed data files from a previously published nTDP dataset of endogenous protein complex previously published by Kelleher and co-workers (MassIVE dataset # MSV000080328) (Skinner *et al*., 2018a). This underlines that MASH Native is capable of analyzing complex nTDP data in the Discovery Mode.

Targeted Mode allows users to comprehensively analyze native top-down or complex-down data for a known protein/protein complex, confirm results generated in Discovery Mode, or potentially find new possible complex associations with database searching. At the MS1 and MS2’ level, MASH Native enables unresolved and isotopically resolved native deconvolution through UniDec (Marty *et al*., 2015). Deconvolution and searching of MS2 or MS3 data in Targeted Mode may be performed using all high-resolution deconvolution algorithms and database search options (*vide supra*). We have used MASH Native to process a native top-down MS dataset of the bovine glutamate dehydrogenase (GDH) hexamer previously published by Loo and co-workers (Li *et al*.,2018) to demonstrated the utility of this targeted workflow (**Figure S7**). MASH Native allowed unresolved MS1 deconvolution and isotopically resolved MS2 deconvolution along with sequence mapping and data visualization in a single software package (**Figure S7**). Recently, our group has demonstrated the utility of MASH for targeted analysis in a complex-down workflow for a native cysteine-linked antibody-drug conjugate (ADC) (**Figure S8**) (Larson *et al*., 2021). The presence of intrachain disulfide bonds limits the fragmentation efficiency of the ADC and reduces sequence coverage by terminal fragment assignment. MASH Native incorporates searching and assignment of internal fragment ions, increasing sequence coverage and revealing sequence coverage of regions bounded by disulfide bonds (**Figure S9**) to provide additional higher-order structural information for proteins and complexes (Lantz *et al*., 2021, 2022).

Since its release on April 7, 2022, MASH Native has been downloaded more than 1,400 times by users all around the world (66 % from North America, 22 % from Europe, 7 % from Asian, Oceania 4%, South America 0.6%, and Africa 0.4%) (**Figure S10**). Noticeably, MASH Native is increasingly gaining popularity and has been well-recognized (Liu *et al*., 2022). Based on the invaluable feedback from the broad community of MASH Users after the first release of MASH Native in Apr. 2022, we have made significant efforts to streamline the installation process and improve user-friendliness of MASH Native in our latest update on December 28, 2022. Additional written and video tutorials have also been added to the MASH website in response to user requests (included in the “Supporting Documents for Users” section of the Supplementary Information), which improves the ease of use and clarifying the best practices for data processing in MASH Native.

## Conclusion

MASH Native provides a unified software solution for the analysis of a variety of complex nTDP data for the first time. As a freely available and universal processing tool, MASH Native is a “one-stop shop” for nTDP data processing that can handle a variety of complex nTDP datasets including unresolved and isotopically MS1, MS2’, MS2, and MS3 in both Discovery and Targeted Modes with database search algorithms as well as data visualization and validation in a user-friendly interface. It can process raw data from various vendor formats and integrates multiple deconvolution/search algorithms into one package. As the nTDP community gains momentum to grow rapidly, MASH Native will play an increasingly important role to streamline nTDP data processing and accelerate the use of nTDP in structural biology and biomedical applications.

## Acknowledgements

This work was supported by the NIH R01 GM125085. We also thank all the MASH users worldwide for their excellent feedback which has helped the development of the software. UniDec development and integration was supported by the National Science Foundation (CHE-1845230 to M.T.M.). J.A.M. acknowledges support from the Training Program in Translational Cardiovascular Science, T32 HL007936-20 and T32 HL007936-21, for funding during the duration of this project. D.S.R. acknowledges the support from the American Heart Association Predoctoral Fellowship Grant No. 832615/David S. Roberts/2021. K.A.B. acknowledges the Vascular Surgery Research Training Program Grant T32 HL110853. K.J.R acknowledges the National Science Foundation Graduate Research Fellowship Program under Grant No. DGE-1747503 and the Graduate School and the Office of the Vice Chancellor for Research and Graduate Education at the University of Wisconsin-Madison, funded by Wisconsin Alumni Research Foundation.

## Notes

### Competing Interest Statement

The authors have declared no competing interest.

### Summary of Updates

The manuscript has been revised to correct a formatting error in the abstract.

## References

Cai, W. et al. (2016) MASH suite pro: A comprehensive software tool for top-down proteomics. Mol. Cell. Proteomics, 15, 703–714.

Catherman, A.D. et al. (2014) Top Down proteomics: Facts and perspectives. Biochem. Biophys. Res. Commun., 445, 683–693.

Chambers, M.C. et al. (2012) A cross-platform toolkit for mass spectrometry and proteomics. Nat. Biotechnol., 30, 918–920.

Chen, B. et al. (2018) Top-Down Proteomics: Ready for Prime Time? Anal. Chem., 90, 110–127.

Cleary, S.P. et al. (2018) Extracting Charge and Mass Information from Highly Congested Mass Spectra Using Fourier-Domain Harmonics. J. Am. Soc. Mass Spectrom., 31–39.

Cleary, S.P. et al. (2016) Fourier Analysis Method for Analyzing Highly Congested Mass Spectra of Ion Populations with Repeated Subunits Sean. Anal. Chem, 88, 6205–6213.

Fellers, R.T. et al. (2015) ProSight Lite: Graphical software to analyze top-down mass spectrometry data. Proteomics, 15, 1235–1238.

Ferrige, A.G. et al. (1991) Maximum entropy deconvolution in electrospray mass spectrometry. Rapid Commun. Mass Spectrom., 5, 374–377.

Horn, D.M. et al. (2000) Automated reduction and interpretation of high resolution electrospray mass spectra of large molecules. J. Am. Soc. Mass Spectrom., 11, 320–332.

Jeong, K. et al. (2020) FLASHDeconv: Ultrafast, High-Quality Feature Deconvolution for Top-Down Proteomics. Cell Syst., 10, 213-218.e6.

Jooß, K. et al. (2022) Native Mass Spectrometry at the Convergence of Structural Biology and Compositional Proteomics. Acc. Chem. Res. 2022, 55, 14, 1928–1937.

Karch, K.R. et al. (2022) Native Mass Spectrometry: Recent Progress and Remaining Challenges. Annu. Rev. Biophys., 51, 157–179.

Keener, J.E. et al. (2021) Native Mass Spectrometry of Membrane Proteins. Anal. Chem., 93, 583–597.

Kou, Q. et al. (2016) TopPIC: A software tool for top-down mass spectrometry-based proteoform identification and characterization. Bioinformatics, 32, 3495–3497.

Lantz, C. et al. (2021) ClipsMS: An Algorithm for Analyzing Internal Fragments Resulting from Top-Down Mass Spectrometry. J. Proteome Res., 20, 1928–1935.

Lantz, C. et al. (2022) Native Top-Down Mass Spectrometry with Collisionally Activated Dissociation Yields Higher-Order Structure Information for Protein Complexes. JACS. 2022, 144, 48, 21826–21830.

Larson, E.J. et al. (2021) High-Throughput Multi-attribute Analysis of Antibody-Drug Conjugates Enabled by Trapped Ion Mobility Spectrometry and Top-Down Mass Spectrometry. Anal. Chem., 93, 10013–10021.

Leney, A.C. and Heck, A.J.R. (2017) Native Mass Spectrometry: What is in the Name? J. Am. Soc. Mass Spectrom., 28, 5–13.

Li, H. et al. (2018) An integrated native mass spectrometry and topdown proteomics method that connects sequence to structure and function of macromolecularcomplexes. Nat. Chem., 10, 139–148.

Liu, R. et al. (2022) Native top-down mass spectrometry for higher-order structural characterization of proteins and complexes. Mass Spectrom. Rev., 1–51.

Liu, X. et al. (2010) Deconvolution and database search of complex tandem mass spectra of intact proteins: A combinatorial approach. Mol. Cell. Proteomics, 9, 2772–2782.

Liu, X. et al. (2012) Protein identification using top-down. Mol. Cell. Proteomics, 11, 1–13.

Loo, J.A. (1997) Studying noncovalent protein complexes by electrospray ionization mass spectrometry. Mass Spectrom. Rev., 16, 1–23.

Marty, M.T. et al. (2015) Bayesian Deconvolution of Mass and Ion Mobility Spectra: From Binary Interactions to Polydisperse Ensembles. Anal. Chem, 87, 4370–4376.

Mcilwain, S.J. et al. (2020) Enhancing Top-Down Proteomics Data Analysis by Combining Deconvolution Results through a Machine Learning Strategy. J. Am. Soc. Mass Spectrom, 2020, 31, 5, 1104–1113.

Melby, J.A. et al. (2021) Novel Strategies to Address the Challenges in Top-Down Proteomics. J. Am. Soc. Mass Spectrom., 32, 1278–1294.

Park, J. et al. (2017) Informed-Proteomics: Open-source software package for top-down proteomics. Nat. Methods, 14, 909–914.

Reid, D.J. et al. (2018) MetaUniDec: High-Throughput Deconvolution of Native Mass Spectra MS Data Set MetaUniDec Deconvolution Integration & Extraction. J. Am. Soc. Mass Spectrom, 30, 118–127.

Sharon, M. and Robinson, C. V (2007) The role of mass spectrometry in structure elucidation of dynamic protein complexes. Annu. Rev. Biochem., 76, 167–193.

Skinner, O.S. et al. (2018a) Multiplexed mass spectrometry of individual ions improves measurement of proteoforms and their complexes. Nat. Chem. Biol., 14, 36–41.

Skinner, O.S. et al. (2018b) Top-down characterization of endogenous protein complexes with native proteomics. Nat. Chem. Biol., 14, 36–41.

Stiving, A.Q. et al. (2019) Surface-Induced Dissociation: An Effective Method for Characterization of Protein Quaternary Structure. Anal. Chem., 91, 190–209.

Sun, R.X. et al. (2016) pTop 1.0: A High-Accuracy and High-Efficiency Search Engine for Intact Protein Identification. Anal. Chem., 88, 3082–3090.

Toby, T.K. et al. (2016) Progress in Top-Down Proteomics and the Analysis of Proteoforms. Annu. Rev. Anal. Chem., 9, 499–519.

Wu, Z. et al. (2020) MASH Explorer: A Universal Software Environment for Top-Down Proteomics. J. Proteome Res., 19, 3867–3876.

Yuan, Z.F. et al. (2012) pParse: A method for accurate determination of monoisotopic peaks in high-resolution mass spectra. Proteomics, 12, 226–235.

Zhou, M. et al. (2020) Higher-order structural characterisation of native proteins and complexes by top-down mass spectrometry. Chem. Sci., 11, 12918–12936.

