## Supplementary Figures and Supporting Documents for "MASH Native: A Unified Solution for Native Top-Down Proteomics Data Processing"

### Supplemental Figures Table of Contents

Figure S1	[S-3](#_Toc6502870)

Figure S2	[S-4](#_Toc6502870)

Figure S3	[S-5](#_Toc6502870)

Figure S4	[S-6](#_Toc6502870)

Figure S5	[S-7](#_Toc6502870)

Figure S6	[S-8](#_Toc6502870)

Figure S7	[S-9](#_Toc6502870)

Figure S8	[S-10](#_Toc6502870)

Figure S9	[S-11](#_Toc6502870)

Figure S10	[S-12](#_Toc6502870)

Supporting Documents for Users	[S-13](#_Toc6502870)

Supplemental References	[S-14](#_Toc6502870)

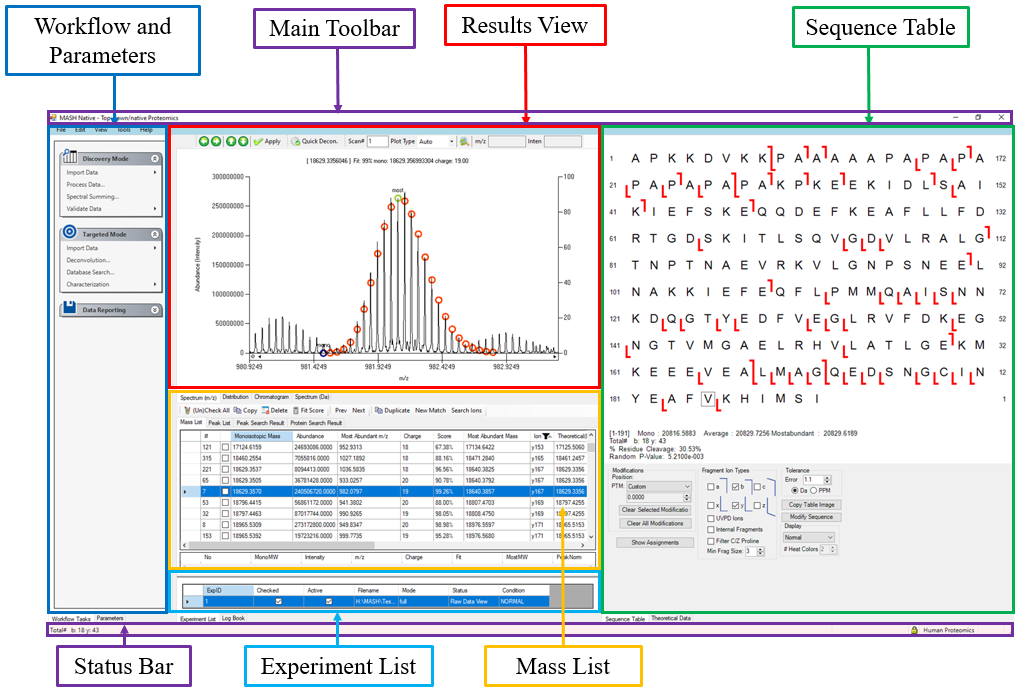

**Figure S1. The main interface of MASH Native includes seven main panels.** 1) The Workflow and Parameters panel handle all the core data processing. Here, users can find the Discovery Mode, Targeted Mode, and Data Reporting Nodes. 2) The Results View panel provides visualization of MS, MS/MS, LC-MS, and LC-MS/MS data. Users can also review spectral deconvolution results, access the Quick Deconvolution feature, and manually adjust the theoretical ion distribution to the actual experimental spectra. 3) The Mass List panel allows users to select deconvoluted fragment ions for manual processing. 4) The Logbook and Status panels provide updates on the progress of data processing. 5) The Experimental List panel allows users to load multiple experiments into MASH Native and easily navigate between them, allowing for efficient processing. 6) The Sequence Table visualizes the fragment ions that match the identified proteoform sequence. 7) The Main Toolbar is where you can exit or minimize the MASH Native window. All these features are discussed in further detail in the MASH Native Supporting Documents, which are automatically downloaded with the software and are also found in the supporting documents linked below (**Supporting Documents 1**–**5**). Video tutorials for new users to MASH Native are linked in the supporting documents (**Supporting Documents 6**–**10**).

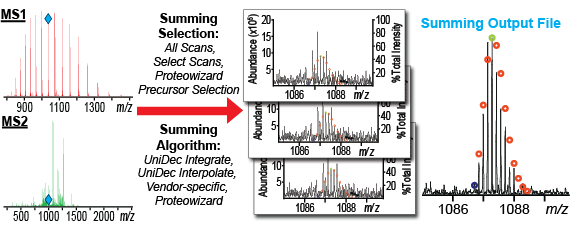

**Figure S2. Spectral summing workflow in MASH Native.** Users can choose to sum all MS2/ MS3 scans, specific scan regions, or use ProteoWizard’s precursor detection algorithms to sum all MS2/MS3 scans for a given precursor ion mass. Once selected, scans can be summed for both MS1 and MS2 using UniDec (Integrate or Interpolate), vendor-specific summing for Thermo data, or ProteoWizard summing to improve signal-to-noise (S/N) (**Supporting Document 4**).

**A**

**B**

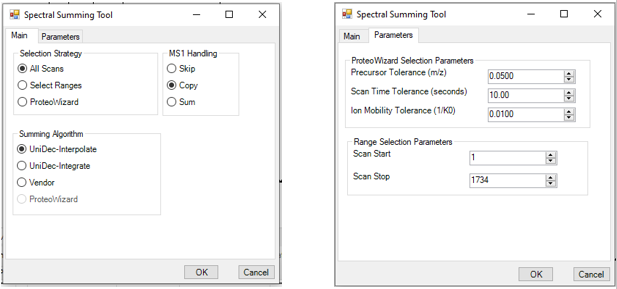

**Figure S3. Spectral Summing Tool in MASH Native.** The spectral summing algorithm allows users to sum scans within their original experiment file. The summing process generates a mzML file which can then be processed by MASH Native, including deconvolution and searching. The summing tool is designed to be flexible by giving users control over the scan selection, the summing algorithm to use, and how to handle MS1 scans in the dataset (A). ProteoWizard selection parameters and scan range selection details can be edited in the “Parameters” tab (B). See the “Best Practices for Spectral Summing” for suggested parameters when performing spectral summing (**Supporting Document 4**).

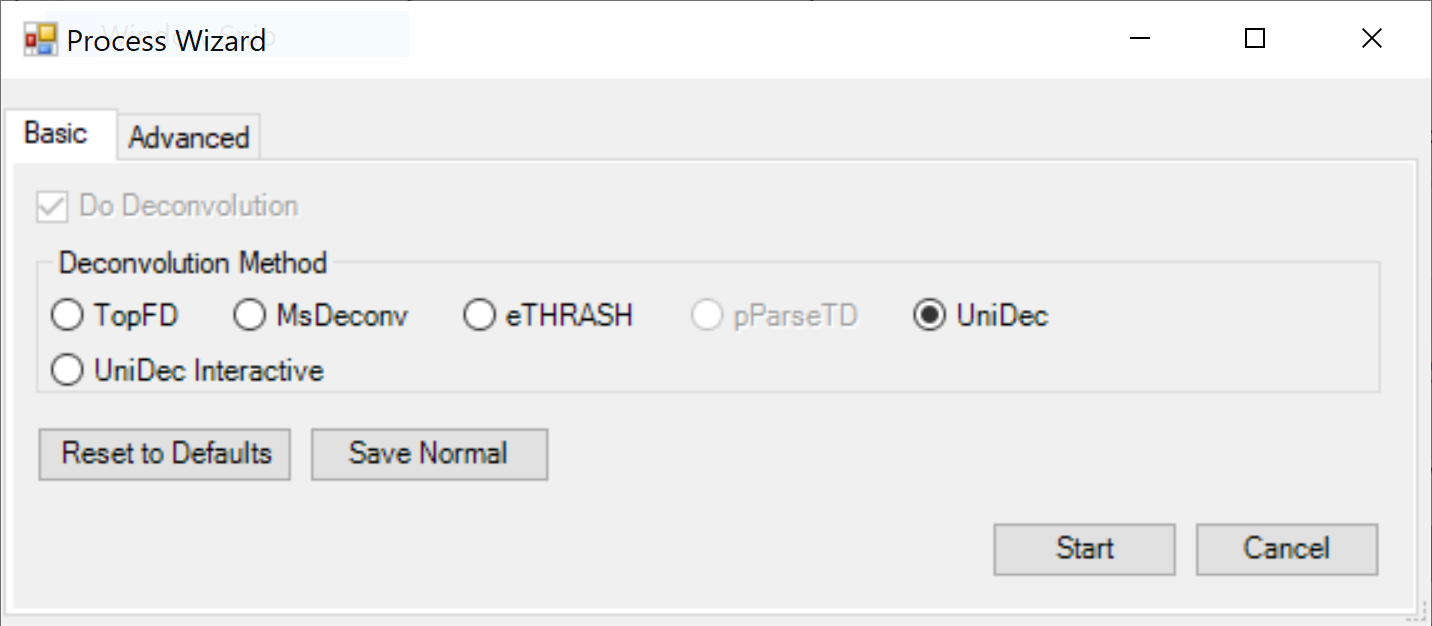

**Figure S4. UniDec deconvolution support in MASH Native.** UniDec deconvolution is a powerful tool to perform charge state deconvolution on both native and denatured protein mass spectra. Correct selection of parameters is critical. The Marty lab has provided a number of different pre-set deconvolution conditions through UniDec which offer parameter suggestions for UniDec for general applications (default), low resolution native MS, high resolution native MS, and isotopically resolved MS (**Supporting Document 5**). Additionally, the Marty lab has recently published an excellent book chapter tutorial to guide user selection of UniDec parameters (Kostelic and Marty, 2022). MASH Native also supports “UniDec Interactive” deconvolution, which allows users load any MASH-compatible data file in the UniDec GUI, perform all processing through the UniDec GUI, then import deconvolution results back into MASH Native for any additional processing.

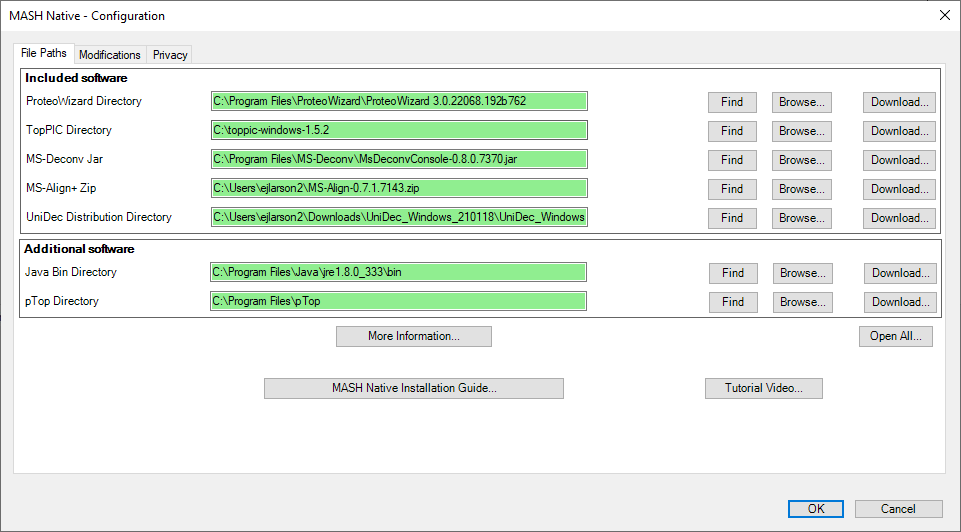

**Figure S5. Configuration of additional deconvolution and search algorithms in MASH Native.** In the MASH Native application, the Configuration tool provides users with an intuitive directory for installation of all associated deconvolution and database search algorithms. In this interface, users can use either the “Find” feature to look for the default directory locations where the software was installed or use “Browse” feature to manually locate the correct directory through a file browser dialog. Clicking the “Download” feature will direct users to the website where the software can be downloaded. Directories found by MASH Native will be displayed in green, while the unidentified directories will be displayed in pink. Software in the “Included software” section is automatically downloaded upon MASH Native installation. “Additional software” is not automatically downloaded but may be installed if users desire (**Supporting Document 2**–**3**). Additionally, a link for the MASH Native Installation Manual can also be found at the bottom of the Configuration tool.

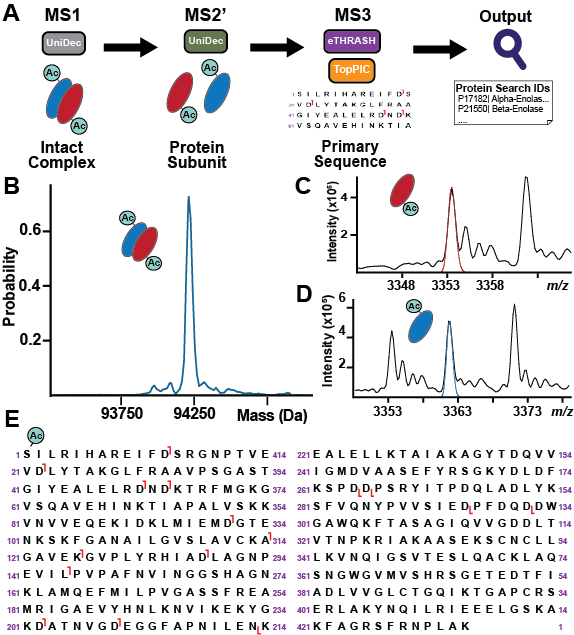

**Figure S6. MASH Native’s Discovery Mode workflow.** Workflow for the analysis of complex-down mass spectrometry data (A). The Discovery Mode in MASH Native was used to process complex-down MS analysis of enolase complex from *mus musculus* (MassIVE dataset # MSV000080328)(Skinner *et al.*, 2018). The Discovery Mode workflow allowed detection of the unresolved intact enolase complex (B), the released α-enolase subunit (C), and β-enolase subunit (D) through low-resolution UniDec deconvolution. High-resolution MS3 processing provided primary sequence coverage for both α-enolase (E) and β-enolase (not shown) and enabled detection and localization of N-terminal acetylation on both subunits. Data can be deconvoluted at the MS1/MS2’ level with either low or high-resolution followed by high-resolution MS2/MS3 fragment spectra deconvolution using a suite of algorithms, including MS-Deconv, pParseTD, eTHRASH, and TopFD. Database search by MS-Align+, pTop, and TopPIC provide the first combination of low-resolution native deconvolution and database search in a single software package.

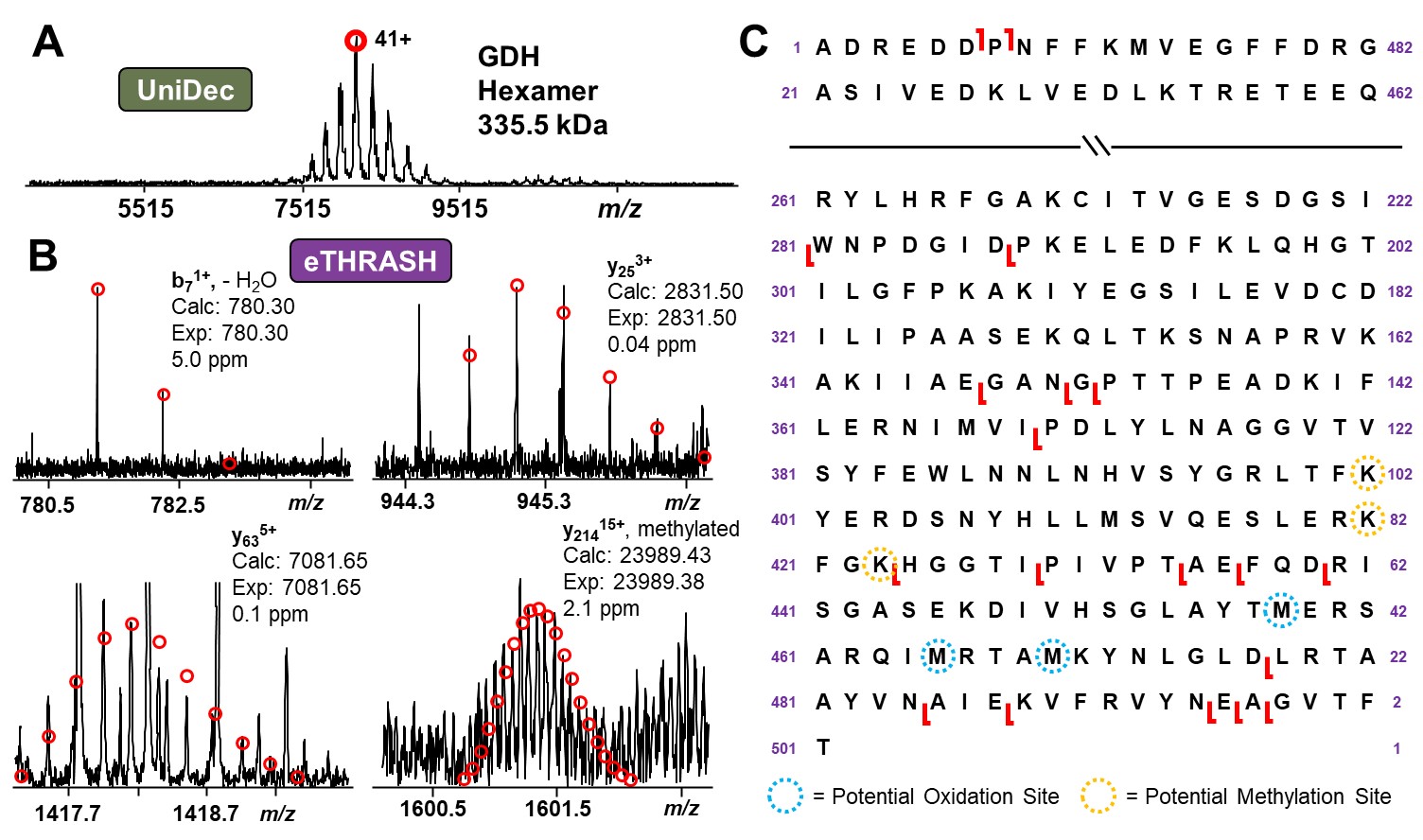

**Figure S7. MASH Native’s Targeted Mode workflow**. Processing of native top-down MS analysis of the hexamer of Bovine glutamate dehydrogenase (GDH) (Li *et al.*, 2018). Analysis of the unresolved MS1 spectra of the GDH hexamer was performed using UniDec (A). Representative fragment ions found using eTHRASH deconvolution of MS2 data (B) and sequence coverage map (C). These results confirm the results reported in Li *et al.* and highlight through MASH Native’s user-friendly and intuitive interface.

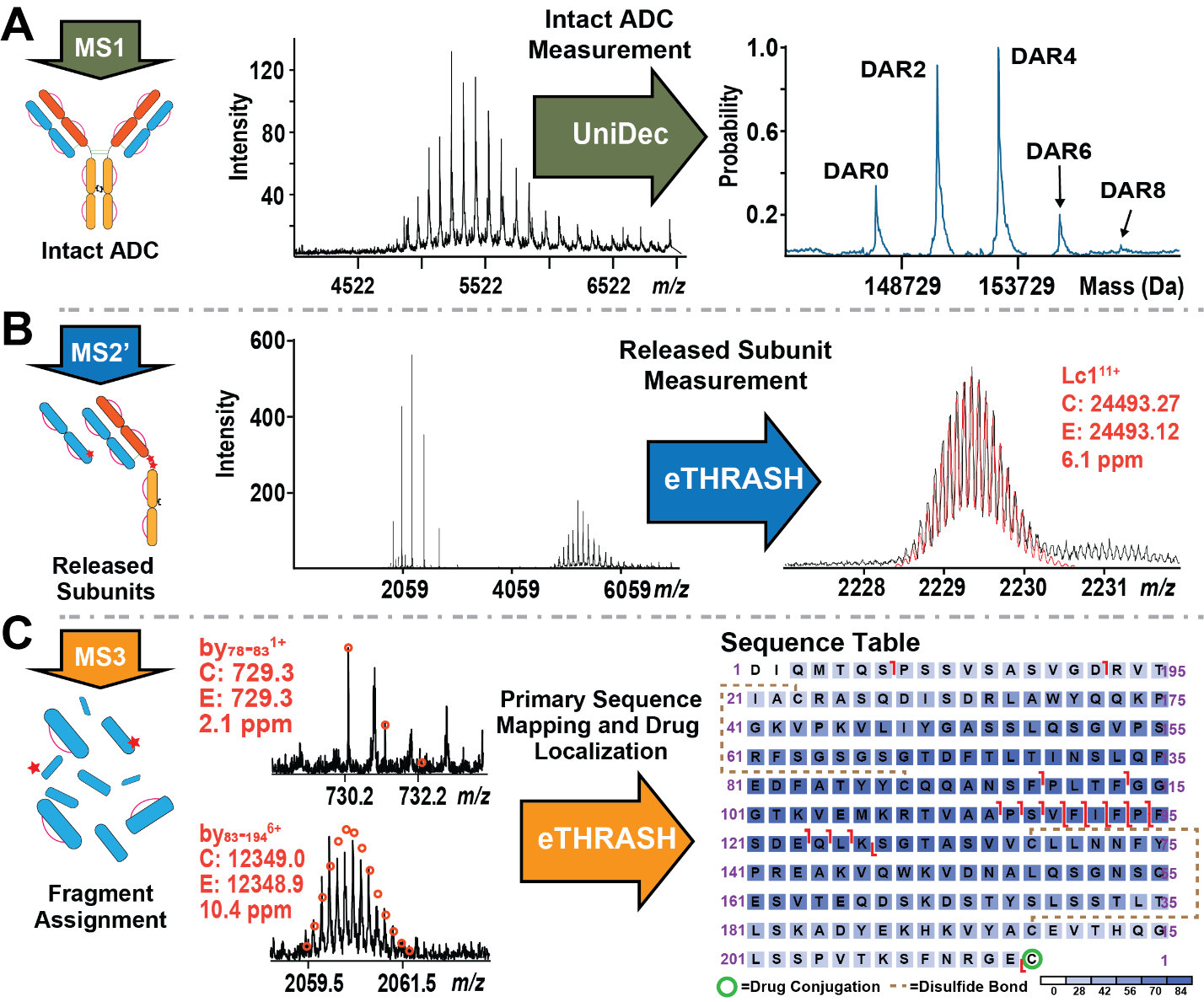

**Figure S8. Analysis of a native cysteine-linked antibody-drug conjugate**. Processing of antibody-drug conjugate (ADC) (Larson *et al.*, 2021). **UniDec processes** unresolved MS1 which provides charge state matching to find individual charge states for specific drug-to antibody ratio (DAR) species and enables viewing of the charge deconvoluted spectra (A). Quantitative output of UniDec even allows calculation of the average DAR value, a critical metric for ADC quality control. High-resolution MS2 deconvolution of collisionally dissociated non-covalently bound species enables high-accuracy subunit mass detection (B). Fragmentation of released subunits, specifically the light chain with one bound drug (Lc1), provides MS3 characterization of the primary sequence through assignment of both terminal and internal fragment ions to confirm the location of intrachain disulfide bonds and the drug binding site (C). Uniquely, internal fragment matches provide sequence coverage in disulfide bound regions, which are not typically accessible to fragmentation by terminal fragment ions.

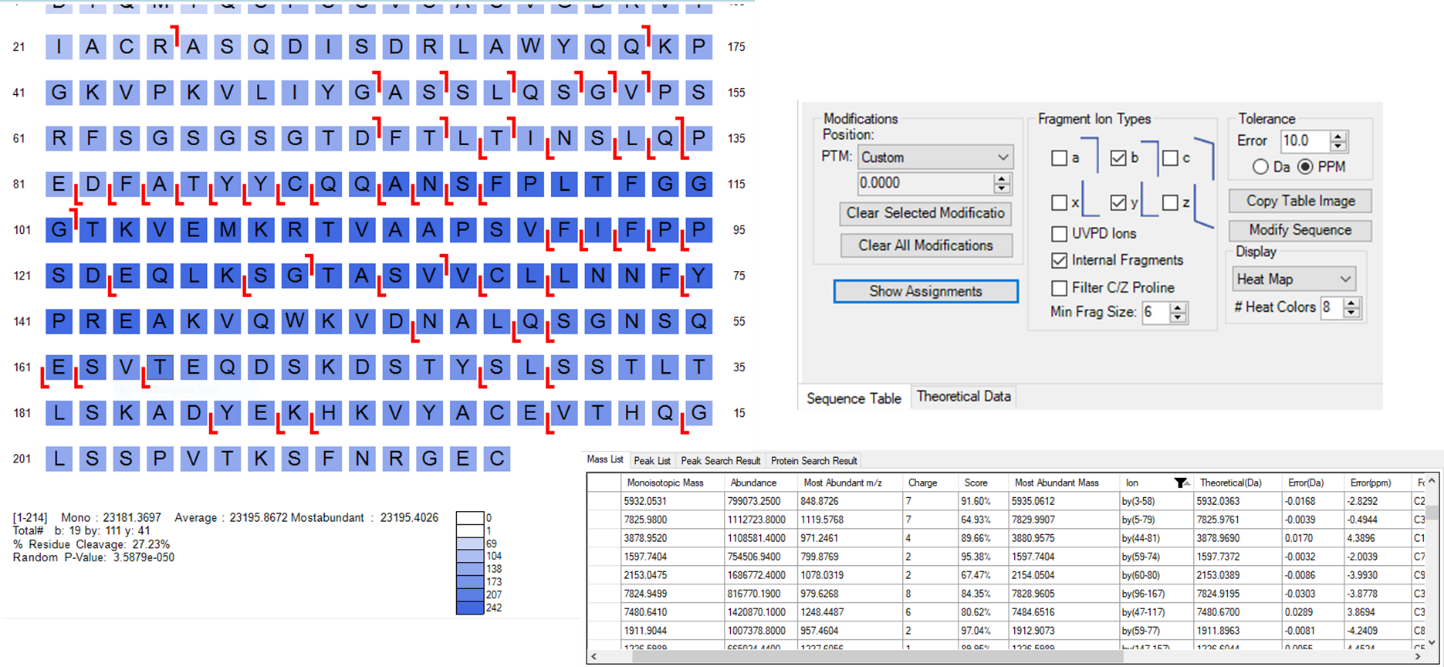

**Figure S9. Internal fragment matching in MASH Native.** In MASH Native’s main interface, users can select “Internal Fragments” under the fragment type and then push the “Show Assignments” function under the sequence table window.

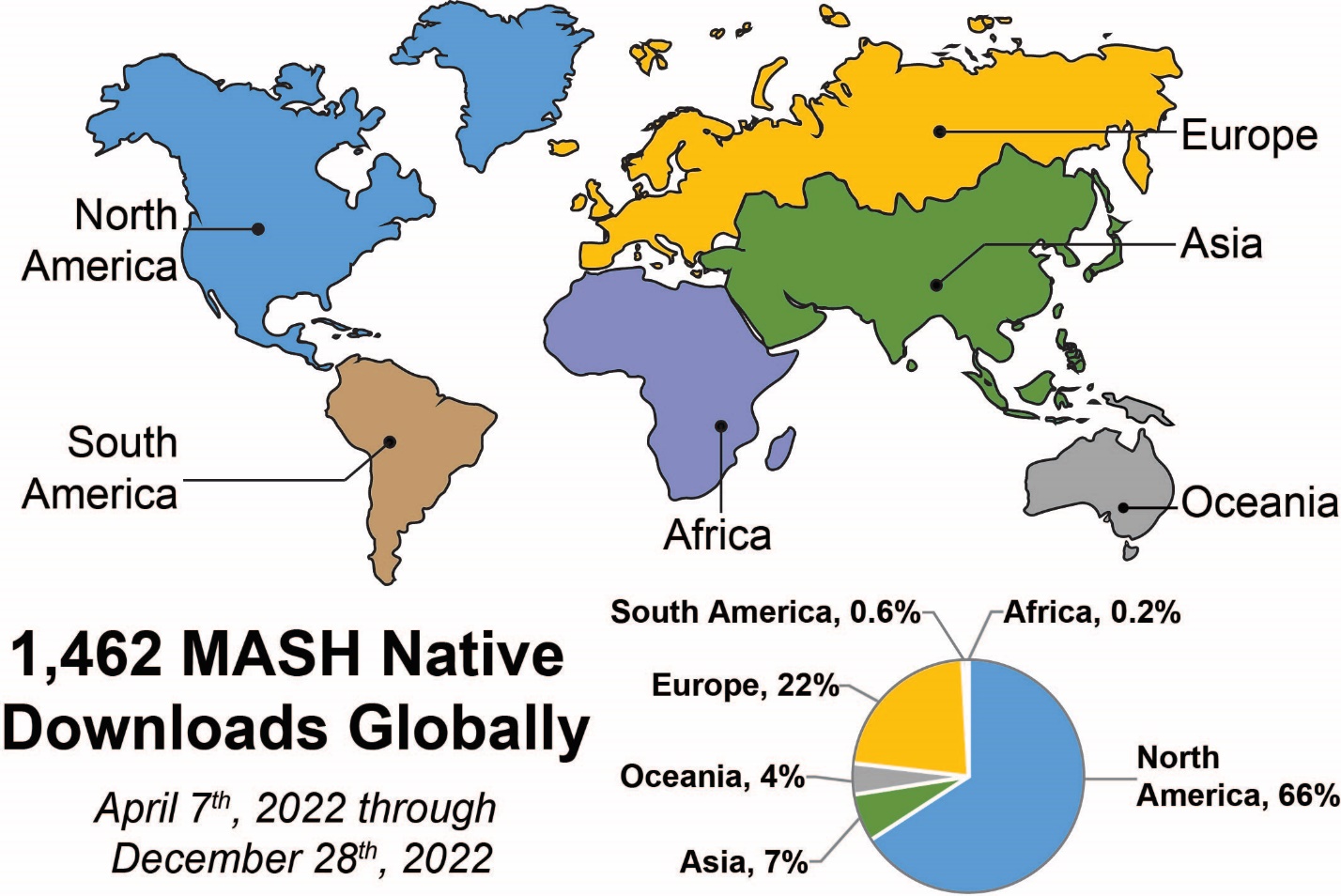

**Figure S10. Global MASH download by geographic region.** MASH Native has been used by many labs globally and is primed to become an integral tool for further developments in native top-down proteomics.

### Supporting Documents for Users

The following MASH Native user documents and video tutorials are provided:

- Supporting document 1 [MASH Native user manual](https://labs.wisc.edu/gelab/MASH_Explorer/doc/Native_UserManual1pt1.pdf)
- Supporting document 2 [MASH Native installation guide](https://labs.wisc.edu/gelab/MASH_Explorer/doc/Native_User%20Installation%20Guide1pt1.pdf)
- Supporting document 3 [MASH Native getting started guide](https://labs.wisc.edu/gelab/MASH_Explorer/doc/Native_GettingStarted1pt1.pdf)
- Supporting document 4 [Best practices for spectral summing](https://labs.wisc.edu/gelab/MASH_Explorer/doc/Best%20Practices%20for%20Spectral%20Summing%20in%20MASH%20Native1pt1.pdf)
- Supporting document 5 [Best practices for UniDec deconvolution in MASH Native](https://labs.wisc.edu/gelab/MASH_Explorer/doc/Best%20Practices%20for%20UniDec%20Deconvolution%20in%20MASH1pt1.pdf)
- Supporting document 6 [Video Tutorial Part 1: Introduction to MASH Native](https://youtu.be/4HmmOjrN9_g)
- Supporting document 7 [Video Tutorial Part 2: MASH Native configuration](https://youtu.be/NtT9q0LDFTM)
- Supporting document 8 [Video Tutorial Part 3: Using the Discovery Mode workflow for identification of an unknown protein](https://youtu.be/hpWGM89fZrw)
- Supporting document 9 [Video Tutorial Part 4: Using the Targeted Mode workflow for characterization of a known protein](https://youtu.be/Qi7TAhLrh4s)
- Supporting document 10 [Video Tutorial Part 5: Post-translational modification analysis using UniDec in MASH Native](https://youtu.be/hjHsrZS2_cI)

**Supplemental References**

Kostelic,M.M. and Marty,M.T. (2022) Deconvolving Native and Intact Protein Mass Spectra with UniDec. *Methods Mol. Biol.*, **2500**, 159–180.

Larson,E.J. *et al.* (2021) High-Throughput Multi-attribute Analysis of Antibody-Drug Conjugates Enabled by Trapped Ion Mobility Spectrometry and Top-Down Mass Spectrometry. *Anal. Chem.*, **93**, 10013–10021.

Li,H. *et al.* (2018) An integrated native mass spectrometry and topdown proteomics method that connects sequence to structure and function of macromolecularcomplexes. *Nat. Chem.*, **10**, 139–148.

Skinner,O.S. *et al.* (2018) Top-down characterization of endogenous protein complexes with native proteomics. *Nat. Chem. Biol.*, **14**, 36–41.
